# Estimating *Pan* evolutionary history from nucleotide site patterns

**DOI:** 10.1101/2022.01.07.475438

**Authors:** Colin M. Brand, Frances J. White, Alan R. Rogers, Timothy H. Webster

**Affiliations:** Bakar Computational Health Sciences Institute, University of California San Francisco, San Francisco, CA, USA; Department of Epidemiology and Biostatistics, University of California San Francisco, San Francisco, CA, USA; Department of Anthropology, University of Oregon, OR, USA; Department of Anthropology, University of Utah, UT, USA

**Keywords:** admixture, bonobos, chimpanzees, Congo River, demography, hybridization

## Abstract

Introgression appears increasingly ubiquitous in the evolutionary history of various taxa, including humans. However, accurately estimating introgression is difficult, particularly when 1) there are many parameters, 2) multiple models fit the data well, and 3) parameters are not simultaneously estimated. Here, we use the software Legofit to investigate the evolutionary history of bonobos (*Pan paniscus*) and chimpanzees (*P. troglodytes*) using whole genome sequences. This approach 1) ignores within-population variation, reducing the number of parameters requiring estimation, 2) allows for model selection, and 3) simultaneously estimates all parameters. We tabulated site patterns from the autosomes of 71 bonobos and chimpanzees representing all five extant *Pan* lineages. We then compared previously proposed demographic models and estimated parameters using a deterministic approach. We further considered sex bias in *Pan* evolutionary history by analyzing the site patterns from the X chromosome. Introgression from bonobos into the ancestor of eastern and central chimpanzees and from western into eastern chimpanzees best explained the autosomal site patterns. This second event was substantial with an estimated 0.21 admixture proportion. Estimates of effective population size and most divergence dates are consistent with previous findings; however, we observe a deeper divergence within chimpanzees at 987 ka. Finally, we identify male-biased reproduction in *Pan* evolutionary history and suggest that western to eastern chimpanzee introgression was driven by western males mating with eastern females.

## Introduction

It is increasingly apparent that hybridization is common, not only in plants but among animals as well (Mallet 2005; Baack and Rieseberg 2007; Hedrick 2013). While some hybridization may be maladaptive, introgressed sequences may also be neutral or even advantageous as well (Baack and Rieseberg 2007; Hedrick 2013). Despite the difficulty in detecting introgression, work using whole-genome sequencing data points to the near ubiquity of introgression in the evolutionary history of large mammals, including bears (Cahill et al. 2015), elephants (Palkopoulou et al. 2018), and hominins (Wall and Hammer 2006; Sankararaman et al. 2016; Vernot et al. 2016; Browning et al. 2018; Slon et al. 2018; Jacobs et al. 2019; Villanea and Schraiber 2019; Rogers et al. 2020). The central role of admixture in hominins suggests that this is likely also true for other non-human primates (Tung and Barreiro 2017).

Our closest living relatives, bonobos (*Pan paniscus*) and chimpanzees (*P. troglodytes*), have been long studied for genomic signatures of admixture. These species can hybridize in captivity (Vervaecke and Van Elsacker 1992), but wild populations are completely separated by the Congo River that may be difficult to traverse. Chimpanzees have been characterized as poor swimmers (Angus 1971) and as being afraid of water (Kano 1992), yet some populations enter bodies of water to forage (Watts et al. 2012a; Watts et al. 2012b) and thermoregulate in hot, dry habitats (Pruetz and Bertolani 2009). Interestingly, bonobos are not characterized as having this aversion to water (Kano 1992) and are known to routinely forage in swamps (Uehara 1990; Hohmann et al. 2019). The geographic distribution of *Pan* prompted early speculation that the formation of this river, which was dated at the time to ∼1.5 - 3.5 Ma, coincided with or prompted speciation in this genus (Horn 1979; Beadle 1981; Myers Thompson 2003), which was estimated to be younger than 1.5 Ma by a number of early genetic studies (Won and Hey 2005; Becquet and Przeworski 2007; Caswell et al. 2008, but see Stone et al. 2002; Yu et al. 2003; Wegmann and Excoffier 2010). Recent work, however, has estimated *Pan* divergence to be older (Prüfer et al. 2012; Prado-Martinez et al. 2013; de Manuel et al. 2016). Further, it now appears that the Congo River is considerably older than previously thought—possibly up to 34 Ma (Takemoto et al. 2015).

The observation of admixture between bonobos and chimpanzees would thus require a connection between the north and south banks of the Congo River unless a *Pan* population crossed the river by swimming and/or rafting or ranged south enough to travel around the headwaters of the Congo River, although the distance makes this second scenario less likely assuming the historical ranges of *Pan* are similar to their current ranges (Takemoto et al. 2015). We note that the impermeability of this geographic barrier is partially a function of river discharge, which can vary widely in both space and time. There is some evidence that river discharge has varied in the recent past, which could create opportunities for both divergence and gene flow (Takemoto et al. 2015), a more plausible scenario given the evidence. Such riverine barriers also separate three of the four chimpanzee subspecies while western chimpanzees occur west of a large forest-savannah mosaic known as the Dahomey Gap (Lester et al. 2021). These rivers also likely experience variation in discharge, which may facilitate introgression between geographically proximate subspecies.

Early analyses for gene flow in *Pan* from autosomal loci yielded inconsistent results and did not include data from Nigeria-Cameroon chimpanzees (Figure 1). Both Hey (2010) and Wegmann and Excoffier (2010) describe gene flow from the common ancestor of chimpanzees into bonobos. Bonobos may have subsequently admixed with different chimpanzee lineages: the ancestor of eastern and central chimpanzees (Wegmann and Excoffier 2010), eastern chimpanzees (Becquet and Przeworski 2007; Wegmann and Excoffier 2010), and central chimpanzees (Wegmann and Excoffier 2010). All studies have found evidence of gene flow among chimpanzee lineages since divergence. Western chimpanzees seem to have admixed with the ancestor of eastern and central chimpanzees and each lineage individually (Won and Hey 2005; Becquet and Przeworski 2007; Caswell et al. 2008; Hey 2010; Wegmann and Excoffier 2010). Three studies also describe eastern-central chimpanzee gene flow (Becquet and Przeworski 2007; Hey 2010; Wegmann and Excoffier 2010). Even with many sites, high-coverage genomic sequences may enable a more robust assessment of *Pan* evolutionary history than multiple autosomal loci.

**Figure 1.**
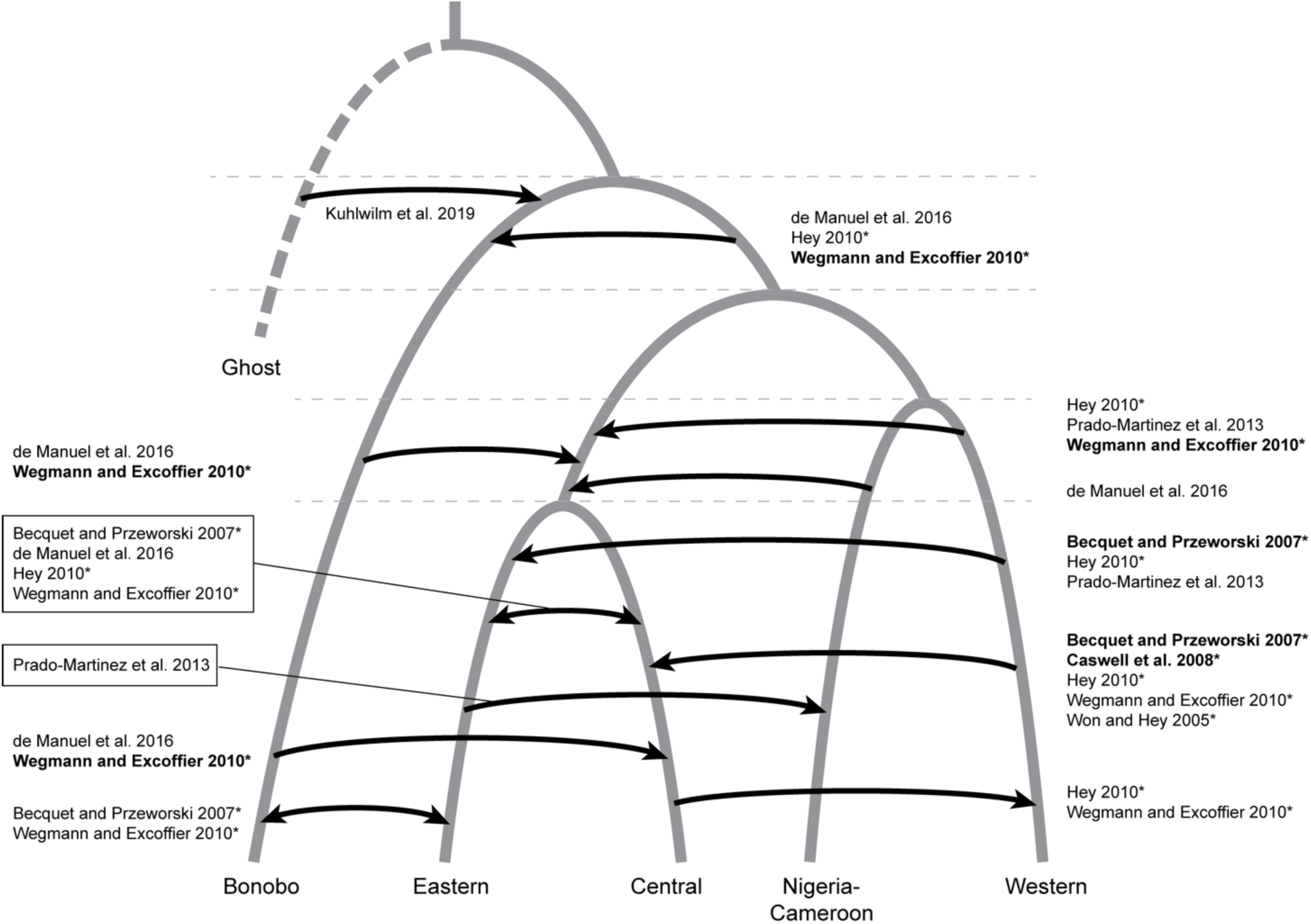
Previous evidence of gene flow in *Pan*. Branch lengths are not to scale. Introgression events only reflect the lineages involved and do not indicate the order/timing of those events between lineage divergences (the dashed horizontal lines). Bolded studies examined the associated event symmetrically rather than asymmetrically. Studies with an asterisk did not consider Nigeria-Cameroon chimpanzees in their analyses.

Admixture inference from whole genome sequences replicated many of these earlier results (Figure 1). Prado-Martinez et al. (2013) was the first whole genome analysis to consider gene flow across all five lineages and noted admixture between eastern and Nigeria-Cameroon chimpanzees, eastern and western chimpanzees, and potential admixture between western chimpanzees and the common ancestor of eastern and central chimpanzees. Given the large number of demographic parameters needed to be estimated under a complex evolutionary history (i.e., many introgression events), de Manuel et al. (2016) estimated parameters using two sets of populations: one that included all lineages except for western chimpanzees and one that included all lineages except for Nigeria-Cameroon chimpanzees. These authors found both additional evidence for bonobo introgression into chimpanzees and evidence for admixture between chimpanzee lineages. The most robust evidence from introgression of bonobos into chimpanzees suggested two events: one that occurred between 200 and 550 ka into the ancestor of eastern and central chimpanzees and a second event < 180 ka, after the eastern and central chimpanzee split (de Manuel et al. 2016). The authors also speculate that the ancestor of common chimpanzees admixed with bonobos deeper in time. Kuhlwilm et al. (2019) further detected introgression from an extinct *Pan* species into bonobos between 377 and 1,627 ka. This ghost lineage was estimated to diverge from the bonobo and chimpanzee common ancestor > 3 Ma. Collectively, these studies point to a complex demographic history, comparable to patterns of introgression observed in humans.

Here, we apply a recently developed method, Legofit (Rogers 2019), to comprehensively compare previously proposed models for *Pan* evolutionary history and estimate demographic parameters: 1) divergence times, 2) effective population sizes, and the 3) timing and degree of introgression. This approach employs site pattern frequencies to infer deep population history by simultaneously estimating all model parameters. There are a few advantages to this approach compared to other commonly used methods for demography. First, within-population variation is ignored and recent changes in population size therefore cannot affect analyses (Rogers 2019). This results in fewer parameters that must be estimated. Second, the uncertainty introduced by statistical identifiability (i.e., when more than one model fits the data well) that is commonly encountered when ascertaining complex demographies can be incorporated into confidence intervals via model averaging (Rogers 2019). Third, simultaneous estimation of all demographic parameters may reduce bias that has been described for other introgression methods (Rogers and Bohlender 2015; Petr et al. 2019). In addition to the autosomes, we also consider the demographic history of the X chromosome in a separate analysis and compare it to the autosomes to assess the potential role of sex bias in *Pan* evolutionary history.

## Methods

### Genomic Data

We used raw short read data on all five extant *Pan* lineages from the Great Ape Genome Project (GAGP) (Prado-Martinez et al. 2013). These sequences represent high coverage genomes from 13 bonobos (*P. paniscus*), 18 central chimpanzees (*P. troglodytes troglodytes*), 19 eastern chimpanzees (*P. t. schweinfurthii*), 10 Nigeria-Cameroon chimpanzees (*P. t. ellioti*), and 11 western chimpanzees (*P. t. verus*). For an outgroup, we also used short read data from a high-coverage human female, HG00513, collected as part of the 1000 Genomes Project (Auton et al. 2015).

### Read Mapping and Variant Calling

We used genotypes generated in Brand et al. (2021). Briefly, these data were reassembled using (a) sex-specific reference genome versions for mapping generated with XYalign (Webster et al. 2019), and (b) a contamination filter during variant calling with GATK4 (Poplin et al. 2018). The use of male- and female-specific versions of the reference genome improves variant calling on the X chromosome (Webster et al. 2019), a critical step for our analyses of sex-bias. The contamination filter was necessary because multiple samples in this dataset suffer from contamination from other samples both within and across taxa (Prado-Martinez et al. 2013). All quality control, read mapping, and variant calling steps are described in detail in Brand et al. (2021) and contained in an automated Snakemake (Köster and Rahmann 2012) available on GitHub (https://github.com/thw17/Pan_reassembly). The repository also contains a Conda environment with all software versions and origins, most of which are available through Bioconda (Grüning et al. 2018).

### Variant Filtration

We excluded unlocalized scaffolds (N = 4), unplaced contigs (N = 4,316), and the mitochondrial genome from these analyses. We used bcftools (Li 2011) to perform further variant filtering and provide command line inputs in parentheses. We first normalized variants by joining biallelic sites and merging indels and SNPs into a single record (“norm -m +any”) using the panTro6 FASTA. We also only included SNPs (“-v snps”) that were biallelic (“-m2 -M2”) and at least 5 bp from an indel (“-g 5”). On a per sample basis within each site, we marked genotypes where sample read depth was less than 10 and/or genotype quality was less than 30 as uncalled (“-S . -i FMT/DP ≥ 10 && FMT/GT ≥ 30”). To ensure that missing data did not bias our results, we further excluded any sites where less than ∼ 80% of individuals (N = 56) were confidently genotyped (“AN ≥ 112”). We also removed any positions that were monomorphic for either the reference or alternate allele (“AC > 0 && AC ≠ AN”). While lack of or low coverage at a locus is problematic, loci with excessive coverage are also of concern. These sites may yield false heterozygotes that are usually the result of copy number variation or paralogous sequences (Li 2014). As our data exhibit a high degree of inter-individual and inter-chromosomal variation in mean coverage (Brand et al. 2021), we applied Li’s (2014) recommendation for a maximum depth filter (d + 4√d, where d is mean depth) to the mean chromosomal coverage of the individual in our sample (*Pan* or *Homo*) with the highest coverage and excluded any loci that exceeded this value (“filter -e FMT/DP > d + 4√d ”) (Table S1). These filtrations steps yielded between 2,413,791,600 and 2,493,198,004 SNVs for our downstream analyses (File S1). After filtration, we generated reference allele frequency (RAF) files for each population that denote the chromosome, the site, the reference allele, the alternate allele, and the frequency of the reference allele.

### Autosomal Analysis

We used Legofit (Rogers 2019; Rogers et al. 2020; Rogers 2021) to estimate demographic history in the five extant lineages of bonobos and chimpanzees. We first used the “sitepat” function (version 1.87) to 1) call ancestral alleles, 2) tabulate site patterns from the RAF files including singletons and 3) generate 50 bootstrap replicates using a moving blocks bootstrap. Ancestral alleles were called using the human genome as an outgroup. We used the allele frequencies within the sample from each population to calculate the probability that a random haploid subsample would exhibit each site pattern. Site pattern labels reflect the samples which exhibit the derived allele; b = bonobo, e = eastern chimpanzee, c = central chimpanzee, n = Nigeria-Cameroon chimpanzee, and w = western chimpanzee. For example, the site pattern cn indicates the central and Nigeria-Cameroon chimpanzee samples have the derived allele at that site.

After visualizing the frequency of the observed site patterns (Figure 2) and examining those for 1) eastern and central chimpanzees (site pattern ec) and 2) Nigeria-Cameroon and western chimpanzees (site pattern nw), we decided to construct two sets of demographic models. In one, eastern and central chimpanzees diverged earlier than Nigeria-Cameroon and western chimpanzees, as possibly suggested by the site patterns. The other set of models considered the reverse and these models are noted by ending in “2”. Next, we constructed various demographic models based on previously proposed introgression events.

**Figure 2.**
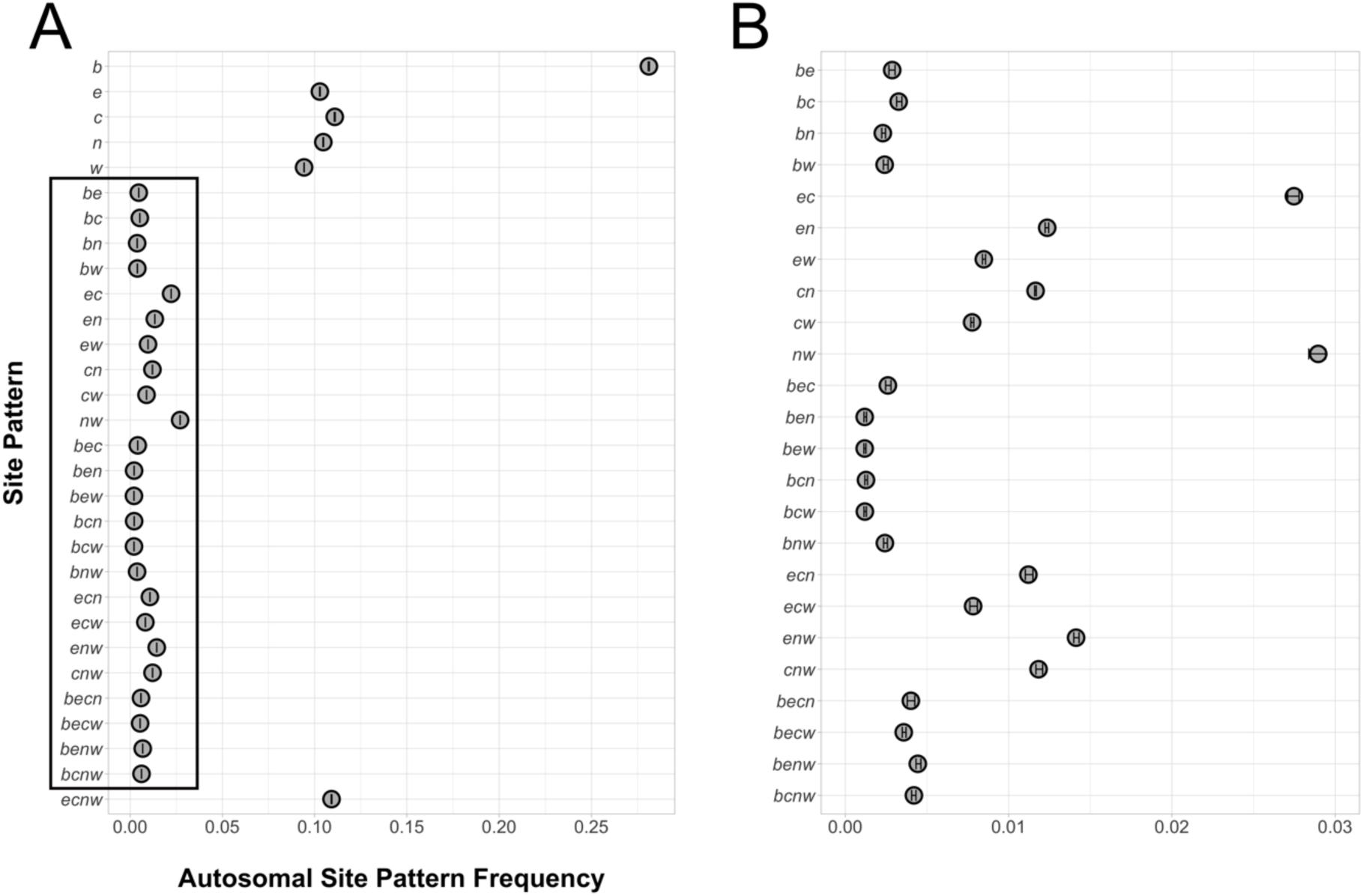
Observed autosomal site patterns. Panel A is the overall distribution of site patterns and Panel B is zoomed in on the region encompassed by the black box on the left. b = *P. paniscus*, e = *P. t. schweinfurthii*, c = *P. t. troglodytes*, n = *P. t. ellioti*, w = *P. t. verus*. Vertical error bars represent the 95% confidence intervals per site pattern.

Legofit cannot easily identify introgression between sister lineages (e.g., between eastern and central chimpanzees after their divergence), so we do not consider those events in this analysis. We prioritized events from whole-genome studies and initially considered all possible subsets of six unidirectional events: α, β, γ, δ, ε, and ζ (N = 64) (Figure 3A). α denotes introgression from a ghost *Pan* lineage into bonobos (Kuhlwilm et al. 2019). β denotes introgression from bonobos into the ancestor of eastern and central chimpanzees (Wegmann and Excoffier 2010; de Manuel et al. 2016). γ denotes introgression from the ancestor Nigeria-Cameroon into eastern chimpanzees (Prado-Martinez et al. 2013). δ denotes introgression from bonobos into central chimpanzees (Wegmann and Excoffier 2010; de Manuel et al. 2016). ε denotes introgression from western into eastern chimpanzees (Becquet and Przeworski 2007; Hey 2010; Prado-Martinez et al. 2013). ζ denotes introgression from western into central chimpanzees (Won and Hey 2005; Becquet and Przeworski 2007; Caswell et al. 2008; Hey 2010; Wegmann and Excoffier 2010). γ consistently resulted in poorly fit models as did introgression from Nigeria-Cameroon chimpanzees into the ancestor of eastern and central chimpanzees (data not shown). Therefore, we excluded this event from the second set of models (N=32) in which eastern and central chimpanzees diverged after Nigeria-Cameroon and western chimpanzees (Figure 3B). We also considered whether adding gene flow from bonobos into eastern chimpanzees (η) and from western chimpanzees into the ancestor of eastern and central chimpanzees (θ) individually and together improved model fit for the five best fitting models (see below). We only considered θ for models that allowed for such an event (i.e., models from Figure 3B).

**Figure 3.**
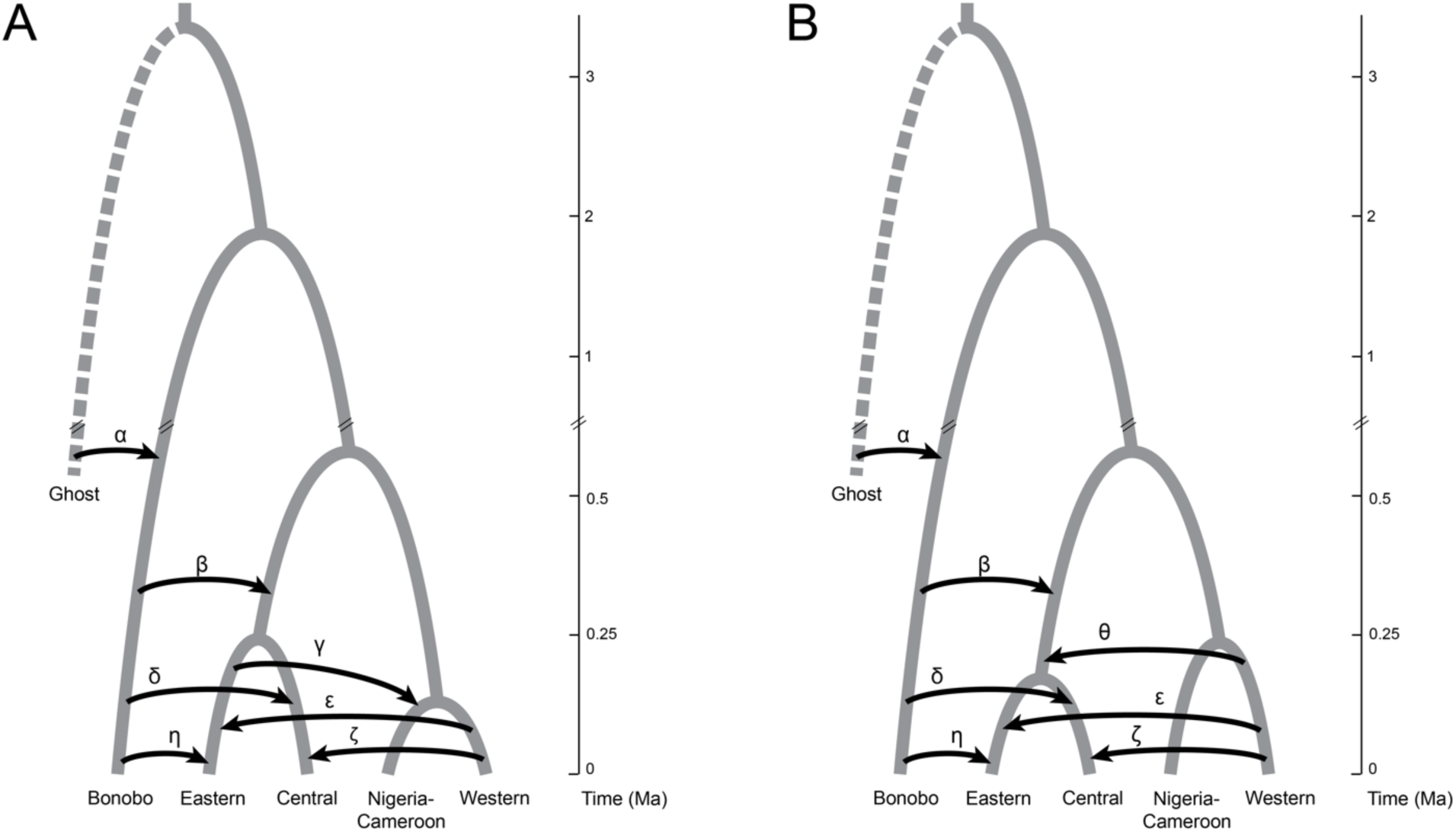
Introgression events considered in this analysis. The the ancestor of eastern and central chimpanzees is older than the ancestor of Nigeria-Cameroon and western chimpanzees based on the observed site patterns. Panel B is a model where these ages are reversed. All divergence times are estimated from de Manuel et al. 2016 and Kuhlwilm et al. 2019. We initially considered all possible subsets of events a,b,c,d,e, and f in Panel A. We then considered all possible subsets all events except for c in Panel B as this event consistently

Legofit requires at least one “fixed” parameter to set the molecular clock, so we chose to set the divergence between bonobos and chimpanzees to the median value as estimated from de Manuel et al. (2016). This value (1.88 Ma) was input in generation units (75,200), based on a generation time of 25 years (Langergraber et al. 2012). While each of the remaining nodes were initially set with the median estimate from de Manuel et al. (2016) (and Kuhlwilm et al. 2019 for models that included α), we designated these parameters to be “free”, which prompts Legofit to generate parameter estimates. We also estimated population size by setting these parameters to be free. We used initial values that ranged from 40,000 diploid individuals for the oldest event to 10,000 individuals for the most recent events. Introgression events were set to be “constrained” parameters, where a parameter is a function of another parameter. Designating parameters as such is useful for reducing the number of free parameters. Events were initially set to occur halfway between one or two divergence parameters or another introgression event. We ordered the timing of the introgression events such that models with multiple introgression events were ordered from oldest to youngest by their Greek letter designation, except for models that included θ, which we placed after β (Figure 3). The order of the more recent chimpanzee introgression events (γ through η) is arbitrary and not based on other results given the discordant findings of previous studies. However, given that these events are more recent and only impact one lineage potentially twice (eastern chimpanzees), we reasoned that event order would not robustly impact model fit. Indeed, this was observed for early analyses (Brand 2021). We did not consider multiple occurrences of the same introgression event. Initial effective population sizes were set to decrease through time such that population sizes decreased upon each divergence. Initial admixture proportions were set to 0.01 for all introgression events (de Manuel et al. 2016), except for ghost admixture into bonobos which was initially set at the median value (0.027) from Kuhlwilm et al. (2019).

Legofit can be run using one of two algorithms: deterministic and stochastic (Rogers 2021). We employed the deterministic algorithm in all models as it is faster and more precise for all but the most complex models (Rogers 2021). We ran the “legofit” function in Legofit (version 2.3.2-3-gd31699a) per demographic model on our real data and each of the 50 bootstrap replicates. Legofit estimates parameters for each model by maximizing the composite likelihood via the “legofit” function. Full likelihood is not maximized because information on linkage disequilibrium is not considered. Legofit employs differential evolution (DE) to maximize composite likelihood. We conducted this in several stages following Rogers et al. (2020). In Stage 1, points in the DE swarm were scattered widely across parameter space and the objective function was evaluated with high precision. As some legofit jobs may converge on different local maxima of the composite likelihood surface, each of the legofit jobs wrote its own swarm of points to a state file. In Stage 2, each legofit job initialized its DE swarm by reading all of the state files produced in Stage 1, enabling legofit to choose among local optima discovered in Stage 1. The evaluation of the objective function was done to very high precision in Stage 2. At this point, we used the “pclgo” function to re-express free variables as principal components. Some free parameters may be tightly correlated, and this can result in broader confidence intervals because there are fewer dimensions than parameters. This issue can be addressed by reducing the dimension of the parameter space. Our early analyses used a value of 0.001 (“--tol 0.001”) such that principal components were only retained if they explained > 0.001% of the variance. However, as the exclusion of dimensions may introduce bias, we retained the full dimension. Re-expression of dimensions as principal components can also improve model fit because it allows legofit to operate on uncorrelated dimensions (Rogers 2021). This step produces a new model file (.lgo file). We then repeated Stages 1 and 2 as Stages 3 and 4 using the new .lgo file.

Models were compared by calculating the bootstrap estimate of predictive error, or bepe, for each model using the “bepe” function in Legofit (Rogers 2019). We also determined whether the top model was superior to all others by implementing the Legofit program “booma.” Briefly, booma calculates weights based on whether each model has the lowest bepe value for the real data and each of the bootstrap replicates (Rogers 2019).

We tested for potential bias in the parameter estimates by generating simulations using msprime (Kelleher et al. 2016) and fitted those simulated data to the model that best fit the observed site patterns. We used parameter point estimates from that model, the previous fixed time parameter (75,200 generations or 1.88 Ma for the *Pan* common ancestor), and used median effective population sizes from Prado-Martinez et al. (2013) for lineages where we did not have an estimate for *N_e_* from our model. We simulated 1 x 10^4^ chromosomes, each 2 x 10^6^ bp in length, and used a mutation rate of 1.5 x 10^-8^ (Besenbacher et al. 2019) and a recombination rate of 1.2 x 10^-8^ (Stevison et al. 2015). This was repeated to generate a total of 50 simulated data sets to which we fit the model using all four stages of the deterministic approach described above. We then visually compared the model’s point estimates to these simulated bootstraps to assess parameter bias.

### X Chromosome Analysis

We used the same methods as described above to generate site patterns for the X chromosome. We used the same methods and criteria to filter SNPs. We further excluded any SNPs from the first pseudoautosomal region (PAR1), the first 2.7 Mb of the X chromosome. PAR1 may exhibit higher genetic diversity than the rest of the chromosomes due to differences in recombination rates, mutation rates, and effective size (Cotter et al. 2016). Because PAR2 and XTR are regions of homology between the X and Y in humans (Charchar et al. 2003; Ross et al. 2005), we used a human female sample as our outgroup to prevent potential biases causes by this homology (Webster et al. 2019). We also generated site patterns for three autosomes of comparable length to the X chromosome: chromosomes 5, 7, and 8. Given that the X chromosome may have a different evolutionary history than the autosomes, we fit each of the previous autosomal models (N=96) to the X chromosome site patterns. As with our autosomal models, we considered whether η and θ improved model fit individually and together for the five best fitting X chromosome models. If the autosomes and X chromosome exhibited the same evolutionary history by sharing the best fitting model, we could assess sex bias in *Pan* demography.

We fit the best autosomal model to chromosome 7 and the X chromosome using the same deterministic approach described above. We input all the estimates parameters as initial values from the model, retained *Pan* divergence as a fixed parameter (1.88 Ma), constrained all the remaining parameters with a free parameter: s, and constrained the admixture proportions with another free parameter: s2. s was set to range from 0 to 10 and given an initial value of 1 for chromosome 7 and 0.75 for the X chromosome. If sex-biased processes are absent from *Pan* evolutionary history, the effective population size inferred from the X chromosome should be 0.75 that inferred from the autosomes (Webster and Wilson Sayres 2016). Thus, departures from 0.75 suggest that female and male effective population sizes were previously unequal. A larger number of breeding males than females should produce s < 0.75, whereas s > 0.75 indicates fewer breeding males than females. s2 was also set to range from 0 to 10 and given an initial value of 1. If migration during introgression was biased toward females, we expected s2 > 1, while s2 < 1 would suggest a greater number of emigrating males.

### Data Availability and Visualization

The raw *Pan* data underlying this article are previously published (Prado-Martinez et al. 2013; de Manuel et al. 2016) and are available from the Sequence Read Archive (PRJNA189439 and SRP018689) and the European Nucleotide Archive (PRJEB15086). The human sample, Biosample ID: SAME123526, is also publicly available from The International Genome Sample Resource website (https://www.internationalgenome.org/data). All models and scripts for this analysis are available on GitHub (https://github.com/brandcm/Pan_Demography). Many figures were generated with R, version 3.6.3 (R Core Team 2020) using ggplot2, version 3.3.3 (Wickham 2016). Correlations between the estimated parameters for the best fit model were visualized in R using corrplot, version 0.90 (Wei and Simko 2021).

## Results

### Introgression from bonobos into the ancestor of eastern and central chimpanzees and from western into eastern chimpanzees best explains Pan nucleotide site patterns

Legofit aligned 2,366,070,805 loci across all six lineages and determined the ancestral allele for 52,809,700 sites. These sites were used to determine site pattern frequencies in the data and 50 bootstrap replicates (Figure 2). As expected, most derived alleles were unique to bonobos, followed by each of the four chimpanzee subspecies and then derived alleles shared across all chimpanzees. Among the remaining non-singleton patterns, the most frequent site patterns were sites unique to Nigeria-Cameroon and western chimpanzees and sites unique to eastern and central chimpanzees. These results are consistent with previously suggested clustering of the chimpanzee subspecies; however, the separation time for Nigeria-Cameroon and western chimpanzees appeared younger than the separation time for eastern and central chimpanzees. Such a pattern could be explained by either 1) eastern and central chimpanzees diverging before Nigeria-Cameroon and western chimpanzees or 2) eastern-central chimpanzees exhibiting a large effective population size. While previous evidence largely supports this second hypothesis (Prado-Martinez et al. 2013; de Manuel et al. 2016), we considered both demographic histories. Hereafter, model names that end with “2” reflect a younger eastern-central divergence time. Each of the remaining site patterns accounted for < 2% of the total distribution.

We ranked an initial set of models by their bepe value encompassing all possible subsets of the α, β, γ, δ, ε, and ζ introgression events (N = 64) and where Nigeria-Cameroon and western chimpanzees diverged more recently than eastern and central chimpanzees (Figure 3a). All models containing γ, or gene flow from eastern chimpanzees into Nigeria-Cameroon chimpanzees (Prado-Martinez et al. 2013), were the lowest ranked models. We also previously considered gene flow from the ancestor of eastern and central chimpanzees into Nigeria-Cameroon chimpanzees (de Manuel et al. 2016) and vice-versa but models containing this event were also consistently the lowest ranked (data not shown). Therefore, we excluded γ when we evaluated models (N = 32) where the ancestor of eastern and central chimpanzees was younger than Nigeria-Cameroon and western chimpanzees (Figure 3b).

We also considered whether gene flow between bonobos and eastern chimpanzees (η) as well as from western into the ancestor of eastern and central chimpanzees (θ) for models that allowed for this scenario (Figure 3b) would improve model fit of our best fit models. We added these events separately and together to the top five models (File S2).

Of these 107 models, we found a single model that best fit the observed site patterns: model βε2 (Table 1). This model includes two episodes of introgression: one from bonobos into the ancestor of eastern and central chimpanzees and a second from western chimpanzees into eastern chimpanzees. This model exhibited small residuals (Figure 4), had the smallest bepe value compared to any other model (1.162 x 10^-5^), and had a weight generated by booma of 1 (File S2); therefore, model averaging was not invoked.

**Figure 4.**
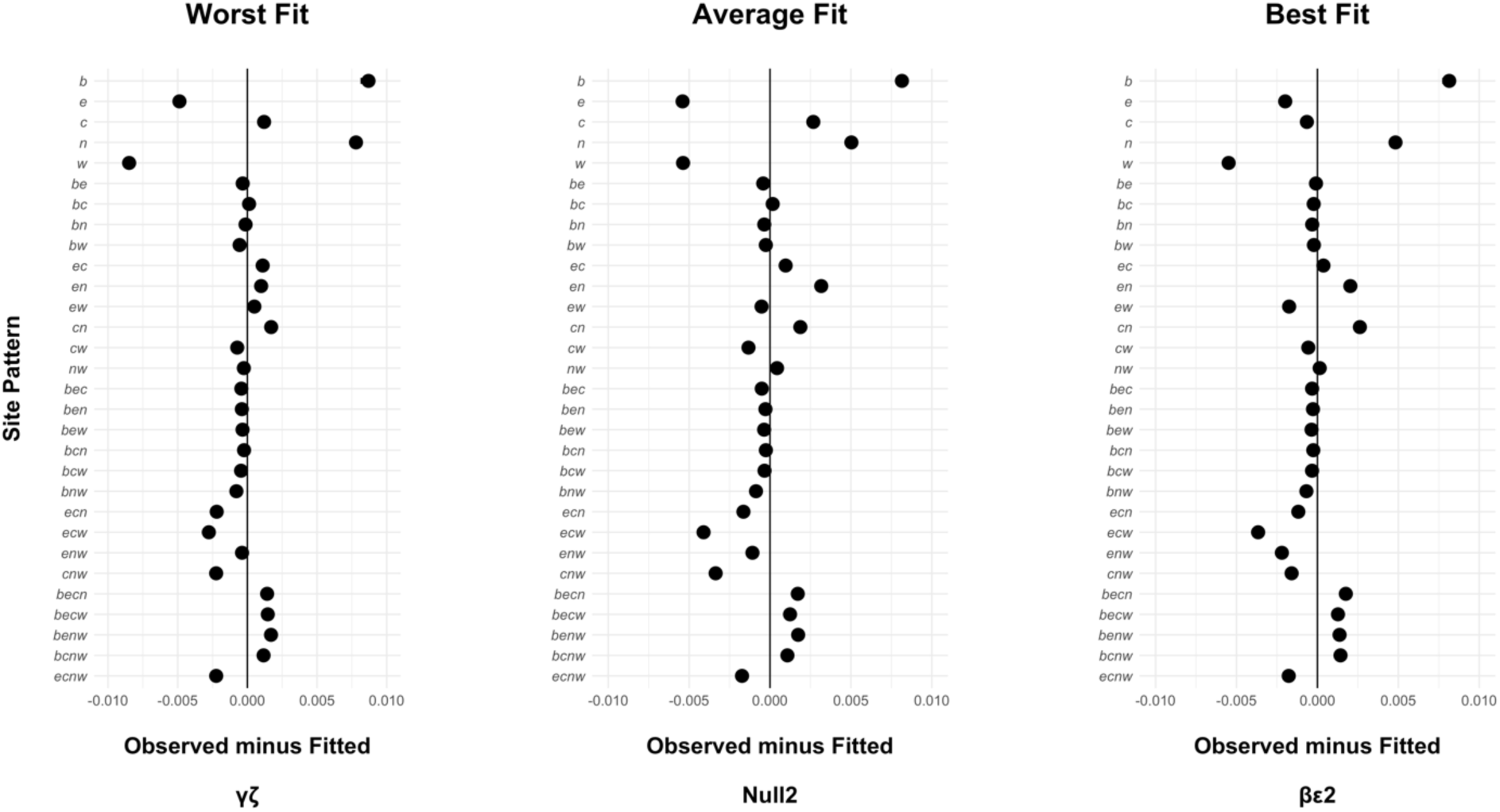
Fitted autosomal model residuals. We display the worst fit model, an average model, and the best fit model. Confidence intervals are plotted but are largely overlaid by the point estimates.

**Table 1.** Best fit autosomal model parameter estimates. ε = introgression from *P. t. verus* into *P. t. schweinfurthii*, β = introgression from *P. paniscus* into the ancestor of *P. t. schweinfurthii* and *P. t. troglodytes*, ec = ancestor of *P. t. schweinfurthii* and *P. t. troglodytes*, nw = ancestor of *P. t. ellioti* and *P. t. verus*, ecnw = common ancestor of all *P. troglodytes* lineages, becnw = *Pan* common ancestor. Admixture is reported as the admixture proportion, population sizes are reported as the number of diploid individuals, and time is reported in years.

|  | Parameter | Point estimate | Lower bound | Upper bound |
| --- | --- | --- | --- | --- |
| Admixture | $\epsilon$ | 0.21402 | 0.20873 | 0.217315 |
| | $\beta$ | 0.00603518 | 0.00539112 | 0.00721139 |
| Population Size | $b$ | 1,107,330 | 96,6960 | 120,1175 |
| | $e$ | 146,408 | 137,731 | 165,923.5 |
| | $w$ | 463,900 | 416,495.5 | 50,1250 |
| | $ec$ | 145,941.5 | 142,575.5 | 149,140.5 |
| | $nw$ | 80,913.5 | 79,786.5 | 82,440 |
| | $ecnw$ | 15,829.6 | 15,741.8 | 15,942.35 |
| | $becnw$ | 37,676.6 | 37,527.3 | 37,955.6 |
| Time | $\epsilon$ | 16,730.125 | 6,707.8 | 30,155 |
| | $\beta$ | 510,447.5 | 500,817.5 | 524,242.5 |
| | $ec$ | 33,460.25 | 13,415.575 | 60,310 |
| | $nw$ | 114,075.5 | 101,862.25 | 121,390.5 |
| | $ecnw$ | 987,437.5 | 985,207.5 | 990,092.5 |

Point estimates and confidence intervals for the βε2 model parameters are provided in Table 1. This model estimated the age for the common ancestor of all chimpanzees to be 987 ka, while the ancestor for western and Nigerian-Cameroon chimpanzees dates to 114 ka and the ancestor of eastern and central chimpanzees was dated to 33 ka. The model also estimated effective population size to vary considerably over time with approximately 38,000 individuals at the time of *Pan* divergence and 16,000 chimpanzees immediately prior to the divergence of the chimpanzee common ancestor. The effective population of both subsequent lineages increased before diverging. The first introgression event in this model occurred from bonobos into the ancestor of eastern and central chimpanzees ∼ 510 ka; however, the estimated admixture proportion was extremely small: 0.006. Given how small this value is, it is possible that the parameter is actually zero and we consider this event to be possible but tentative. The second introgression event is estimated to have occurred 16 ka and suggests ∼ 21% of the eastern chimpanzee genome is derived from western chimpanzees.

After simulating data using this model, calculating site patterns, and fitting these site patterns to the model, we found minimal bias in our parameter estimates for admixture and the effective population size of older events (Figure 5). The effective population size for the individual lineages appears to be underestimated despite the high estimates for these parameters. It is possible that these values are an artifact our of approach. Both the ancestor of eastern and central chimpanzees and the ancestor of Nigeria-Cameroon and western chimpanzees appear to be underestimated as well. Point estimates for younger divergence times appear to be overestimates such that the extant chimpanzee subspecies may be younger than we calculate here. This also appears to be true for the timing of the introgression events themselves. However, our estimated age for the common ancestor of all chimpanzees agreed with the simulated data.

**Figure 5.**
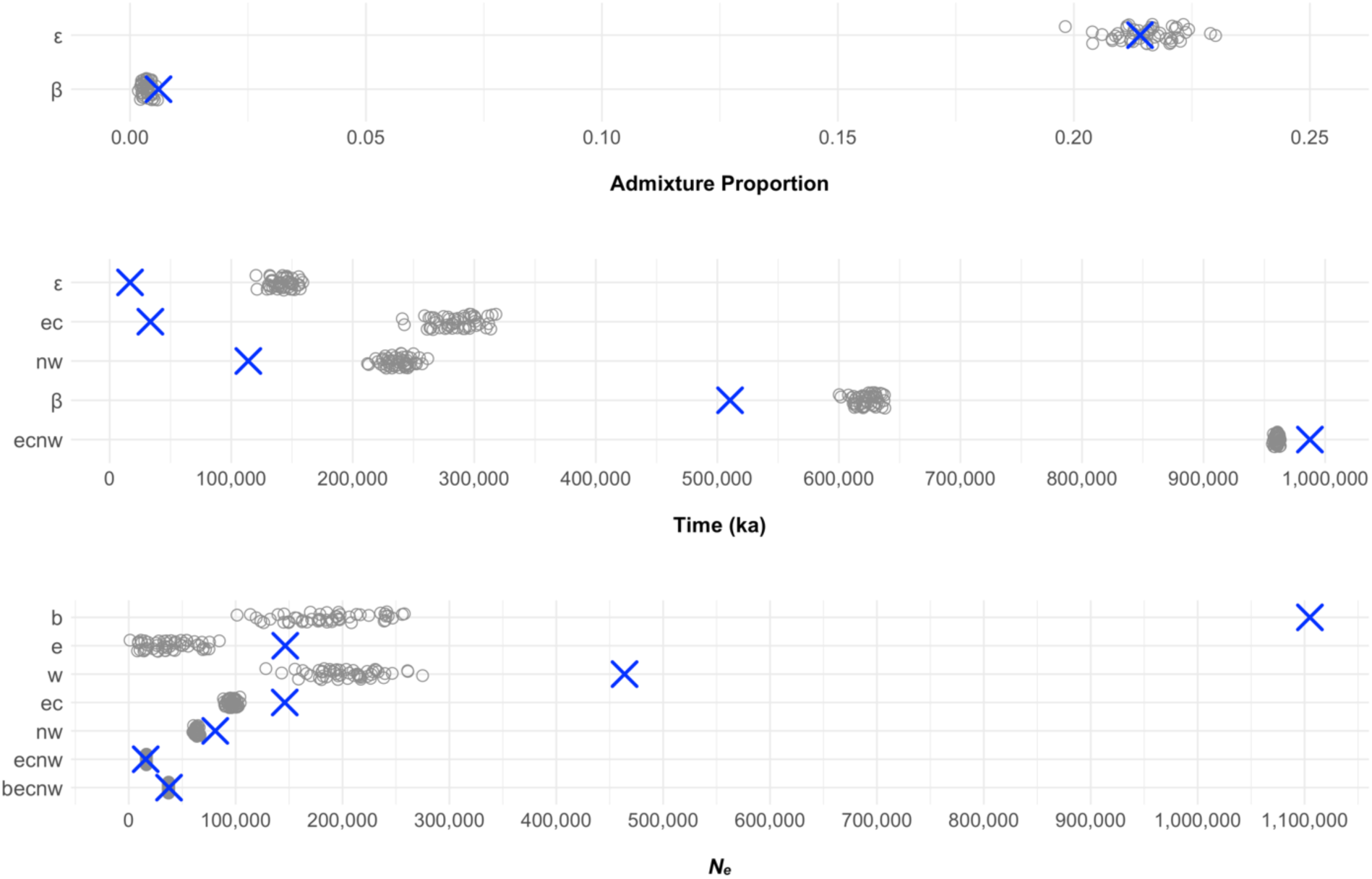
Parameter estimate bias. The orange points represent point estimates for the parameters from the βε2 model. Open gray circles represent 50 values estimated by legofit using site patterns generated from data simulated with the δ model parameters using msprime. If the simulated data are < the point estimate, the point estimate is underestimated, while if the simulated data are > the point estimate, the point estimate is overestimated. ε = introgression from *P. t. verus* into *P. t. schweinfurthii*, β = introgression from *P. paniscus* into the ancestor of *P. t. schweinfurthii* and *P. t. troglodytes*, ec = ancestor of *P. t. schweinfurthii* and *P. t. troglodytes*, nw = ancestor of *P. t. ellioti* and *P. t. verus*, ecnw = common ancestor of all *P. troglodytes* lineages, becnw = *Pan* common ancestor.

**Figure 6.**
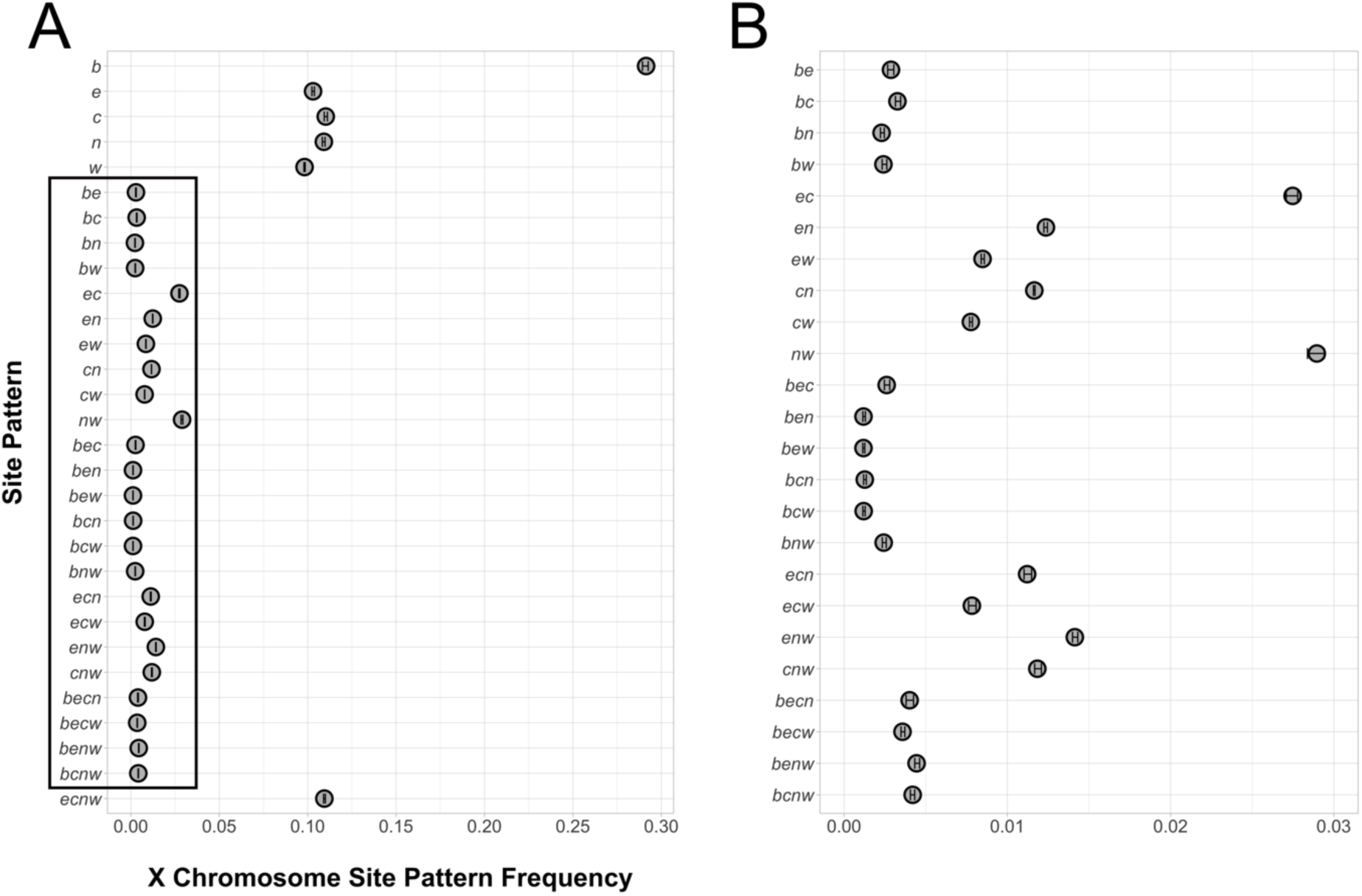
Observed X chromosome site patterns. Panel A is the overall distribution of site patterns and Panel B is zoomed in on the region encompassed by the black box on the left. b = *P. paniscus*, e = *P. t. schweinfurthii*, c = *P. t. troglodytes*, n = *P. t. ellioti*, w = *P. t. verus*. Vertical error bars represent the 95% confidence intervals per site pattern.

### X chromosome site patterns are also explained by the autosomal introgression events and potentially additional gene flow

Next, we considered the added complexity of sex bias in *Pan* evolutionary history. We calculated site patterns for the X chromosome and three similarly sized autosomes: chromosomes 5, 7, and 8. Site patterns could be determined for 1,932,892 loci on the X chromosome, or 3.7% of the loci used in the autosomal analyses, and exhibited similar patterns to chromosomes 5, 7, and 8 (Figure 7). First, we fit these site patterns to the best autosomal model, βε2. All four chromosomes had nearly identical residuals with the X chromosome exhibiting larger confidence intervals for most of the site patterns (Figure 7). Despite this fit, we proceeded to evaluate all our previous autosomal models (N = 96) as the X chromosome may have a different evolutionary history than the autosomes. As with the autosomal analysis, we considered whether bonobo into eastern chimpanzee (η) and western chimpanzee into eastern and central chimpanzee introgression (θ) improved the fit of our top five X chromosome models individually and together. Three models exhibited low bepe values: βεη2, βε2, and αβδεηθ2 (File S3). These models were comparable such that the top model was not superior to the others. This resulted in two models that were differentially weighted: βεη2 = 0.863, βε2 = 0, and αβδεηθ2 = 0.137 (File S3).

**Figure 7.**
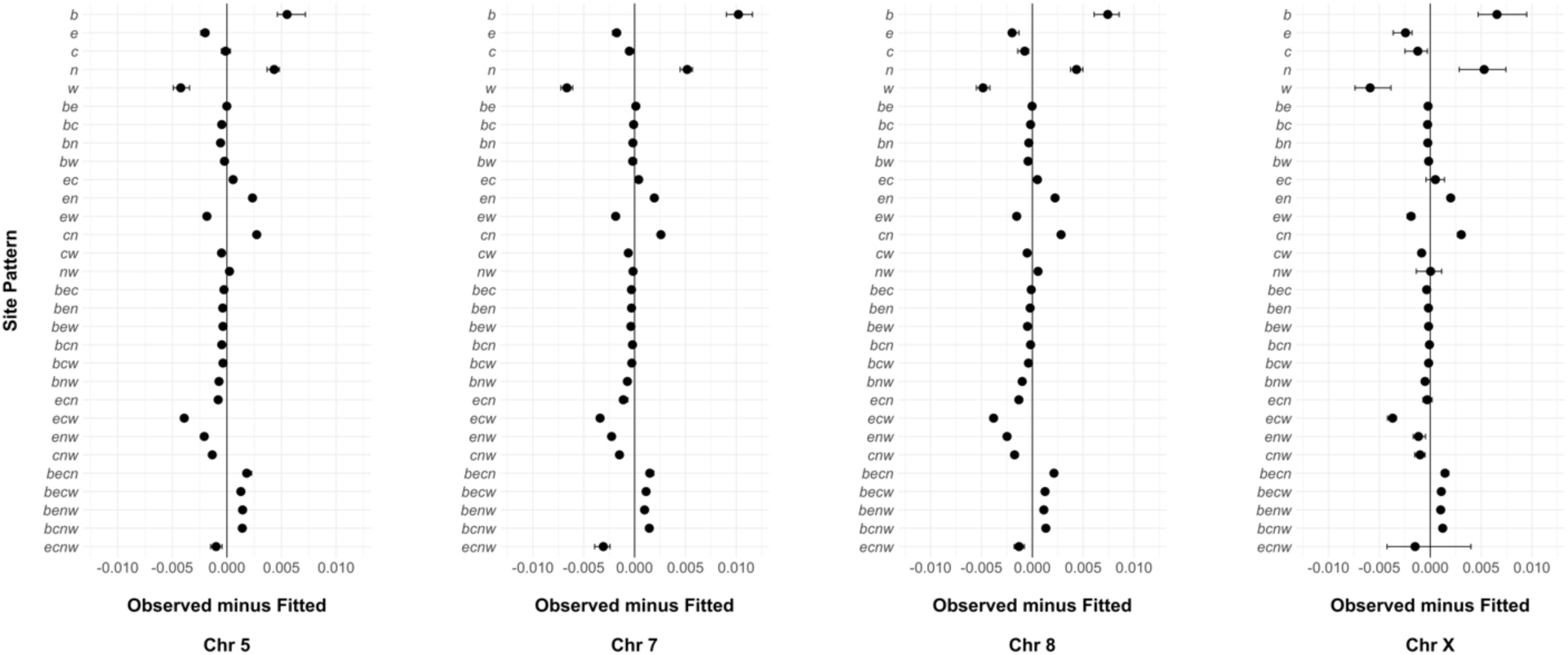
Fitted individual autosomes and X chromosome model residuals. All models were fit to the best autosomal model: βε2. Confidence intervals are plotted but are largely overlaid by the point estimates.

Table 2 summarizes model averaged parameters for the X chromosome top model set. Most effective population size and time parameters were congruent with the best autosomal model. The X chromosome model also included the α introgression event (introgression of an extinct Pan lineage into bonobos), which was estimated to be 0.07, and yielded a population size of 178,530 and divergence time of 2.03 Ma for bonobos, chimpanzees, and this extinct *Pan* lineage. In addition to α, the averaged model included introgression from bonobos into central chimpanzees (δ), bonobos into eastern chimpanzees (η), and western chimpanzees into the ancestor of eastern and central chimpanzees (θ). Admixture proportions for these first two events ranged from small (δ = 0.019) to negligible (η = 0.008) and reversing the direction of η did not improve model fit (data not shown). However, admixture proportion of the older chimpanzee introgression event, θ, was comparable to that of the western into eastern chimpanzee event: 0.186. Similar to the best autosomal model, the population size estimates for introgressing lineages were unreasonably large. As with the autosomes this may be an artifact of our approach.

**Table 2.**
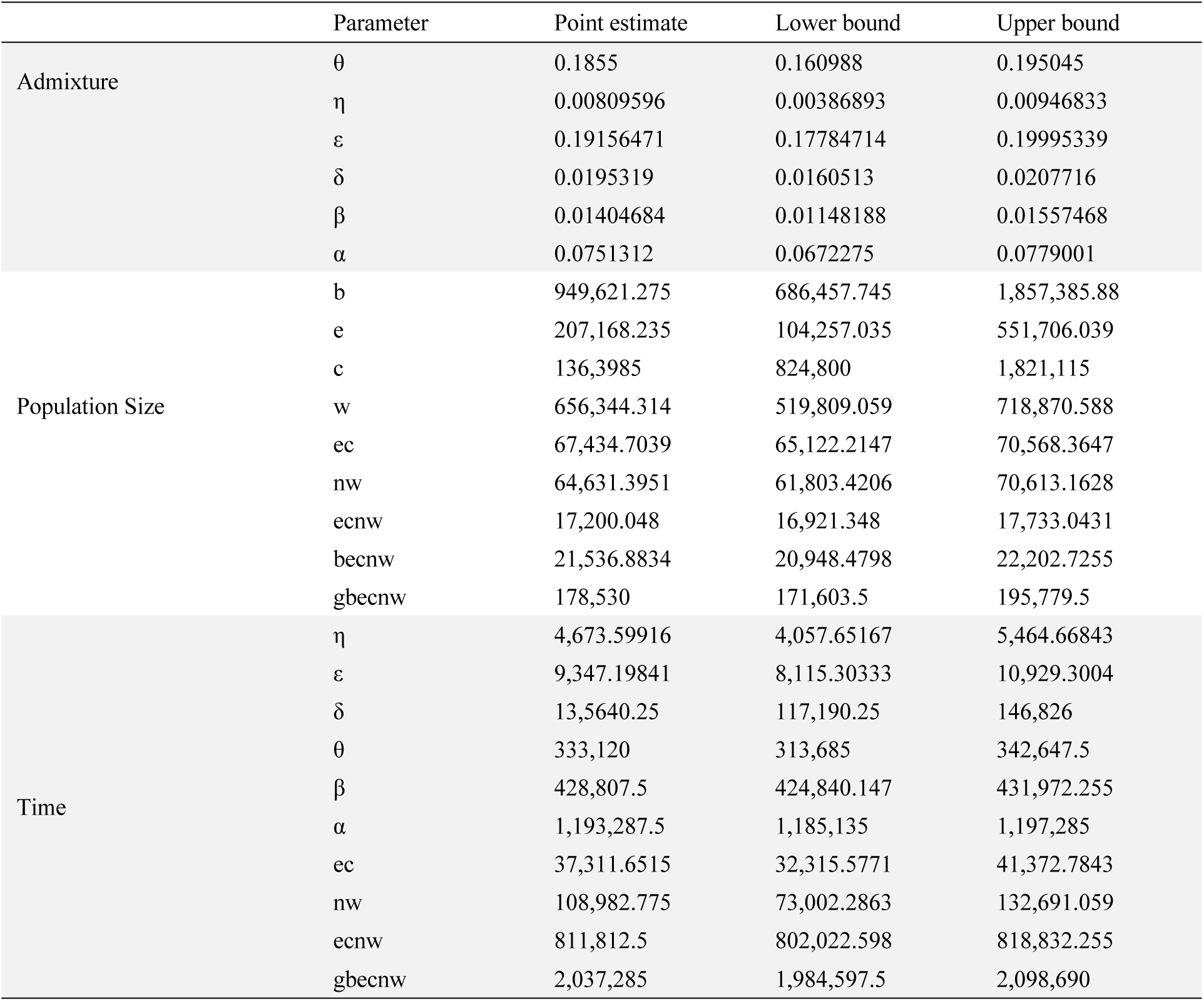
Model averaged X chromosome parameter estimates. θ = introgression from *P. t. verus* into the ancestor of *P. t. schweinfurthii* and *P. t. troglodytes*, η = introgression from *P. paniscus* to *P. t. schweinfurthii*, ε = introgression from *P. t. verus* into *P. t. schweinfurthii*, δ = introgression from *P. paniscus* into *P. t. troglodytes*, β = introgression from *P. paniscus* into the ancestor of *P. t. schweinfurthii* and *P. t. troglodytes*, α = introgression from an extinct Pan lineage into *P. paniscus*, ec = ancestor of *P. t. schweinfurthii* and *P. t. troglodytes*, nw = ancestor of *P. t. ellioti* and *P. t. verus*, ecnw = common ancestor of all *P. troglodytes* lineages, becnw = *P. paniscus* and *P. troglodytes* common ancestor, gbecnw = *Pan* common ancestor. Admixture is reported as the admixture proportion, population sizes are reported as the number of diploid individuals, and time is reported in years.

|  | Parameter | Point estimate | Lower bound | Upper bound |
| --- | --- | --- | --- | --- |
| Admixture | $\theta$ | 0.1855 | 0.160988 | 0.195045 |
| | $\eta$ | 0.00809596 | 0.00386893 | 0.00946833 |
| | $\varepsilon$ | 0.19156471 | 0.17784714 | 0.19995339 |
| | $\delta$ | 0.0195319 | 0.0160513 | 0.0207716 |
| | $\beta$ | 0.01404684 | 0.01148188 | 0.01557468 |
| | $\alpha$ | 0.0751312 | 0.0672275 | 0.0779001 |
| Population Size | $b$ | 949,621.275 | 686,457.745 | 1,857,385.88 |
| | $e$ | 207,168.235 | 104,257.035 | 551,706.039 |
| | $c$ | 136,3985 | 824,800 | 1,821,115 |
| | $w$ | 656,344.314 | 519,809.059 | 718,870.588 |
| | $ec$ | 67,434.7039 | 65,122.2147 | 70,568.3647 |
| | $nw$ | 64,631.3951 | 61,803.4206 | 70,613.1628 |
| | $ecnw$ | 17,200.048 | 16,921.348 | 17,733.0431 |
| | $becnw$ | 21,536.8834 | 20,948.4798 | 22,202.7255 |
| | $gbecnw$ | 178,530 | 171,603.5 | 195,779.5 |
| Time | $\eta$ | 4,673.59916 | 4,057.65167 | 5,464.66843 |
| | $\varepsilon$ | 9,347.19841 | 8,115.30333 | 10,929.3004 |
| | $\delta$ | 13,5640.25 | 117,190.25 | 146,826 |
| | $\theta$ | 333,120 | 313,685 | 342,647.5 |
| | $\beta$ | 428,807.5 | 424,840.147 | 431,972.255 |
| | $\alpha$ | 1,193,287.5 | 1,185,135 | 1,197,285 |
| | $ec$ | 37,311.6515 | 32,315.5771 | 41,372.7843 |
| | $nw$ | 108,982.775 | 73,002.2863 | 132,691.059 |
| | $ecnw$ | 811,812.5 | 802,022.598 | 818,832.255 |
| | $gbecnw$ | 2,037,285 | 1,984,597.5 | 2,098,690 |

### *Pan* evolutionary history is characterized by male-biased reproduction and introgression

While the site patterns from the X chromosome support a slightly different evolutionary history than the autosomes, this history includes both introgression events estimated for the autosomes. Indeed, the top-ranked autosomal model was ranked second for the X chromosome based on its bepe value and a model with a single additional introgression event comprises the majority of the weight on the averaged X chromosome model. Therefore, we decided to estimate historic differences in female and male effective population sizes and migration rates using the best autosomal model on the X chromosome and a similarly sized autosome: chromosome 7. The estimates for the chromosome 7 multipliers agreed with our predictions. We found that the chromosome 7 effective population size and time parameters scaled closely to those from all autosomes, s = 0.990964, and the admixture proportions were nearly identical: s2 = 1.01159. All else equal, the hemizygosity of the X chromosome in males should result in population size and time parameters from X chromosome site patterns that scale by 0.75, while sex biases will drive that value up or down depending on both the measure and direction of sex bias (Webster and Wilson Sayres 2016). We estimated s to be 0.92512, indicating more breeding females than breeding males. Further, s2 was 0.317951, which suggests male-biased admixture. While the β event is questionable, this s2 multiplier points to interbreeding occurring between western chimpanzee males and eastern chimpanzee females more frequently than vice-versa.

## Discussion

Studies have used various approaches to estimate demographic parameters for *Pan* evolutionary history. These estimates vary in their agreement with one another and there is no clear agreement on the number and distribution of introgression events in this lineage (Figure 1). In this study, we not only estimate parameters but also comprehensively compare previous models. We find that a model (βε2) containing two introgression events best fits *Pan* autosomal nucleotide site patterns: 1) bonobo introgression into the ancestor of eastern and central chimpanzees and 2) western chimpanzee introgression into eastern chimpanzees. The admixture proportion of this first event, bonobos into chimpanzees, was estimated to be 0.006. This small value suggests that this event may not have even occurred, which would mean an even simpler evolutionary history. Despite the uncertainty surrounding this event, the other event appears to have involved substantial admixture and both events have important implications for *Pan* biogeography.

We estimate that approximately 21% of eastern chimpanzee DNA is derived from western chimpanzees. Our simulations from the best fitting model that generated this parameter indicate that this admixture proportion is unlikely to be biased. This is also true for the tentative gene flow event from bonobos to the ancestor of eastern and central chimpanzees, which has been previously described (Wegmann and Excoffier 2010; de Manuel et al. 2016). This gene flow event would have likely possible given variation in the discharge of the Congo River and such contact could have happened at many points along that river and its tributaries. Indeed, sections of the Congo River near Kisangani appear to be strong candidates for such a location based on current hydrology (Takemoto et al. 2015). Evidence of gene flow from bonobos into the ancestor of eastern and central chimpanzees is further evidenced by the possible adaptiveness of putatively introgressed bonobo alleles in chimpanzees (Nye et al. 2018).

Admixture from western into eastern chimpanzees, who presently occupy the ends of the species’ geographic range, is perplexing when examined based on current biogeography. However, such an event is easily explained by differences in the current and historic range of these taxa. Variation in suitable chimpanzee habitat, including that for eastern and western chimpanzees, is well described for the past 120 ka, particularly forest refugia (e.g., Barratt et al. 2021). Contact between these lineages would require a connection through or north of the Dahomey Gap and may have occurred northeast of the current Nigeria-Cameroon chimpanzee range, possibly in Cameroon, Central African Republic, Chad, or Nigeria. The exact history of the Dahomey Gap is only partially understood but multiple primate species occur on both sides of the gap in the Upper Guinean and Congolian (or Lower Guinean) rainforests (Harcourt and Wood 2012). Further, paleoenvironmental data suggest the Dahomey Gap has been subject to fluctuating periods of forest cover since at least 1.05 Ma (Dupont et al. 2001). Some of the periods may have resulted in substantial forest expansion (Dupont and Weinelt 1996; Miller and Gosling 2014), enough to potentially allow for the introgression event supported by this study. Further, it’s clear that chimpanzees occurred outside their present range at least once in the deep past based on the recovery of fossil teeth from the Kapthurin Formation in Kenya (McBrearty and Jablonski 2005).

The timing of western chimpanzee introgression into eastern chimpanzees is unclear. Our estimate of the times of divergence for the ancestor of eastern and central and the ancestor of Nigeria-Cameroon and western chimpanzees are much more recent than expected (Becquet and Przeworski 2007; Hey 2010; Prado-Martinez et al. 2013; de Manuel et al. 2016), and our assessment of parameter bias suggests this might even be overestimated. This would point to a very recent divergence for eastern and central chimpanzees, < 30 ka, implying that the introgression from western chimpanzees occurred within the past few thousand years. While possible, it seems more plausible that these lineages diverged around the times proposed by other studies, ∼ 100 and 250 ka. Admixture following divergence, as evidenced by broad time parameter confidence intervals, may lead Legofit to infer a more recent point estimate. A more tractable approach to dating the western into eastern introgression event would involve the identification of putatively introgressed loci or haplotypes and assessing their age.

Our estimate of the age of the common chimpanzee (*P. troglodytes*) ancestor (∼ 987 ka) appears to be robust and is consistent with expectations from simulations of the best fitting autosomal model. However, our estimate is hundreds of thousands of years older than other estimates (e.g., 544 - 633 ka (de Manuel et al. 2016)). We note that this estimate is largely consistent across the 107 autosomal models evaluated. Further, the estimate for this parameter from X chromosome site patterns (∼ 812 ka) is similar.

Our estimates for population size largely support previous findings (Prado-Martinez et al. 2013; de Manuel et al. 2016). Following divergence, the common ancestor of all chimpanzees experienced a period of decline. This was followed by substantial increases in both the ancestor of Nigeria-Cameroon and western chimpanzees and particularly the ancestor of eastern and central chimpanzees. The estimated *N_e_* for each lineage suggests that each subspecies experienced a population decline after divergence with their common ancestor. However, our population size estimates for each lineage at the time of introgression is puzzling. We found a large effective population size for bonobos, eastern chimpanzees, and western chimpanzees. This may represent an instance of statistical identifiability where parameters are correlated, resulting in a broader confidence interval (Rogers 2019). Indeed, some of these parameters are tightly correlated with each other and the beta admixture proportion (Figure S1). Such a high parameter values could also be explained by geographic population structure (Nei and Takahata 1993). Bonobo population structure has been inferred from cranio-dental morphology (Pilbrow and Groves 2013), malarial infection (Liu et al. 2017), and mitochondrial haplotypes (Kawamoto et al. 2013, but see Eriksson et al. 2004). The geographic origins for the bonobos used in this analysis are unknown and the eastern and western chimpanzees used here also span a large geographic range that could result in population structure (Prado-Martinez et al. 2013). Population structure would result in increased effective population sizes and warrants further study. Another potential explanation for large effective population sizes is gene flow between bonobos and an extinct sister lineage; a different lineage than the previously proposed ghost lineage.

While our best model for demographic history was similar to that of the autosomes, there were some interesting differences. The best model (βεη2) had an additional introgression event from bonobos to eastern chimpanzees. Further, a more complex model (αβδεηθ2) was weighted to ∼ 0.13 and included an additional three introgression events; one of which, western chimpanzees into the ancestor of eastern and central chimpanzees exhibited a substantial admixture proportion: ∼ 0.18. This reflects that the X chromosome may capture additional facets of *Pan* evolutionary history due its size, inheritance patterns, and hemizygosity in males. However, one might predict reduced introgression of autosomal events, rather than additional events, based on patterns of reduced Denisovan and Neanderthal ancestry in human X chromosomes, regions described as “introgression deserts” (Sankararaman et al. 2016; Vernot et al. 2016). The support for ghost admixture in our X chromosome model is particularly perplexing because the original proposal for this event found that the bonobo X chromosome was largely devoid of ghost *Pan* ancestry. This result warrants further investigation, but one major difference between studies is the correction for sex-chromosome mismapping in this study. Specifically, regions of homology on the sex chromosomes lead to read mismapping and downstream technical artifacts variant calls (Webster et al. 2019). Thus, our correction might have increased our power for detecting this ghost admixture on the X chromosome. We also draw attention to the reduced power that is inherent to studying the X chromosome; however, we feel that statistical power is not an issue here as the results of the X chromosome closely match similarly-sized autosomes.

Despite exhibiting an equal sex ratio among adult individuals, sex-biased reproduction in *Pan* is well described (Gerloff et al. 1999; Boesch et al. 2006; Inoue et al. 2008; Wroblewski et al. 2009; Newton-Fisher et al. 2010). Indeed, extended periods of sexual receptivity in bonobos as compared to chimpanzees has prompted the hypothesis that male competition for mating opportunities is reduced in bonobos compared to chimpanzees (Hare et al. 2012), resulting in lower reproductive skew. However, the bonobo communities studied to date at LuiKotale and Wamba, exhibit higher reproductive skew than all but one eastern chimpanzee community (Surbeck et al. 2017; Ishizuka et al. 2018; McCarthy et al. 2020). While we cannot speak to differences between bonobos and chimpanzees, we found that the estimated population size and time parameters did not scale as expected compared to similarly sized autosome, providing evidence of male reproductive skew throughout *Pan* evolutionary history. Comparison of admixture proportions between the X chromosome and chromosome 7 also suggests sex-bias in the introgression events described in this study, such that western chimpanzee males mated with eastern chimpanzee females more than vice versa. This scenario is intriguing because western chimpanzee males are slightly larger than eastern chimpanzee males (Smith and Jungers 1997) and therefore may be more likely to win in disputes over females. We caution that reduced estimates of admixture on the X chromosome compared to the autosomes may also be the result of purifying selection, which is expected to be more efficient on the X chromosome due to its hemizygosity in males as proposed for the absence of archaic ancestry in humans (Sankararaman et al. 2016; Vernot et al. 2016).

There are several important considerations for this analysis. First, Legofit is unable to estimate subsequent gene flow between recently diverged lineages; therefore, other introgression events may have occurred that we cannot directly model using this approach. However, if gene flow occurred shortly after a lineage diverged, we would expect the confidence interval this time parameter to be quite large. This may be the case for the ancestors of both eastern and central chimpanzees whose lower and upper bound estimates span a considerable time period. Yet, this interval is small for the divergence of Nigeria-Cameroon and western chimpanzees (range: ∼ 20 ka) and even smaller for the common ancestor of all chimpanzees (range: ∼ 5 ka). As we set the bonobo-chimpanzee divergence time as the fixed parameter in this analysis, we do not have a resulting confidence interval to infer subsequent gene flow as estimated by other studies. Therefore, in addition to gene flow from bonobos into the ancestor of eastern and central chimpanzees and from western into eastern chimpanzees, admixture between bonobos and chimpanzees and between eastern and central chimpanzees may also have occurred.

We note that the parameters estimated from this analysis were generated by setting one fixed parameter (the *Pan* divergence date or Tbecnw) to set the molecular clock. However, the point estimate used in this analysis was the median of a range from de Manuel et al. (2016). Thus, if the true divergence date is different to that used here, our parameter estimates would change as well. Additional genomic data from bonobos and chimpanzees may yield more accurate estimates of this critical parameter. The ordering of events may influence parameter estimates beyond the timing of each introgression event as well as model fit although this seems unlikely. We also did not allow for an introgression event to occur multiple times (e.g., multiple pulses of introgression between two lineages). A better approach for determining multiple events is estimating the age of introgressed regions (e.g., Plagnol and Wall 2006; Sankararaman et al. 2014; Hubisz et al. 2020). Differently aged haplotypes in the same lineage would point to multiple events (de Manuel et al. 2016) and we encourage further study of this in future research.

The site patterns of derived alleles in bonobos and chimpanzees confirm multiple aspects of their evolutionary history while offering new insights into other facets. We find support for one introgression event from western into eastern chimpanzees. However, the biogeography of this event remains difficult to explain without invoking differences in the range of these subspecies over the course of the late Pleistocene compared to the present. Collectively, the best fit demographic model is simpler than more recently proposed models. Finally, our results point to a deeper divergence time for common chimpanzees. Additional genomic and paleoenvironmental data would be immensely informative in deciphering the evolutionary history of our closest living relatives and may provide insight into the evolution of other taxa in this region during this time period, including humans.

## Acknowledgements

The ideas and analyses presented here benefited from helpful discussion with many individuals. We thank current and former members of the Sterner and Ting labs: Tanner Anderson, Savanah Bird, Diana Christie, Elisabeth Goldman, Enrique Gomez, Claire Goodfellow, Alex Hickmott, Jenneca McCarter, Sam Queeno, Noah Simons, Kirstin Sterner, Jessica Stone, Nelson Ting, and Hannah Wellman. We also thank members of the Capra lab for their feedback: Tony Capra, Shiron Drusinsky, Sarah Fong, Erin Gilbertson, Bian Li, Evonne McArthur, Souhrid Mukherjee, David Rinker, and Keila Velázquez-Arcelay. This work was funded by the National Science Foundation (NSF BCS 1945782). We are grateful for computational resources and support from the University of Utah Center for High Performance Computing and access to the University of Oregon computing cluster, Talapas.

