## Supplement for "Estimating *Pan* evolutionary history from nucleotide site patterns"

Figure S1. Correlations between parameter estimates using the observed data and 50 bootstrap replicates. Parameters that begin with “T” refer to time whereas parameters that begin with “twoN” refer to effective population size. “beta” and “epsilon” are the estimated admixture proportions and “Tbeta” and “Tepsilon” are the time estimates for those events. b = *P. paniscus* at the time of introgression, c = *P. t. troglodytes* at the time of introgression, ec = ancestor of *P. t. schweinfurthii* and *P. t. troglodytes*, nw = ancestor of *P. t. ellioti* and *P. t. verus*, ecnw = common ancestor of all *P. troglodytes* lineages, becnw = *Pan* common ancestor.

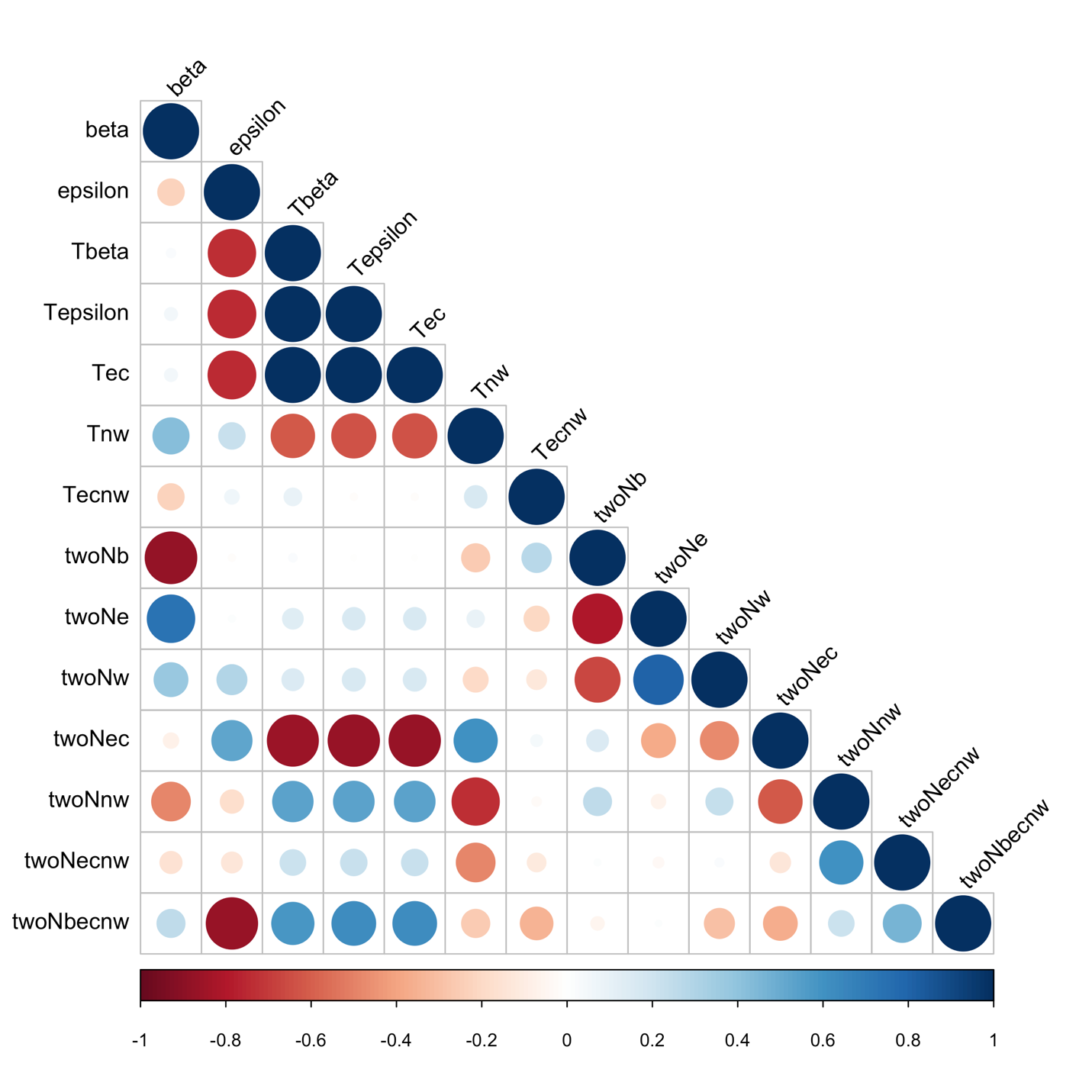

Table S1. Max depth criterion for filtering SNPs per chromosome. The cutoff was calculated as d + 4√d where d is the mean read depth for that chromosome. We used the highest read depth, which varied by chromosome between the *H. sapiens* individual and a *P. t. schweinfurthii* individual (Cleo).

| Chromosome | Cleo | HG00513 | Max Depth Cutoff |
| --- | --- | --- | --- |
| chr1 | 57.18 | 61.16 | 92.4 |
| chr10 | 58.01 | 60.86 | 92.1 |
| chr11 | 57.44 | 61.86 | 93.3 |
| chr12 | 56.27 | 62.97 | 94.7 |
| chr13 | 53.4 | 67.29 | 100.1 |
| chr14 | 56.56 | 63.16 | 94.9 |
| chr15 | 57.99 | 60.91 | 92.1 |
| chr16 | 59.77 | 52.04 | 90.7 |
| chr17 | 62.84 | 52.82 | 94.5 |
| chr18 | 55.45 | 64.83 | 97.0 |
| chr19 | 64.04 | 44.28 | 96.0 |
| chr20 | 66.88 | 58.01 | 99.6 |
| chr21 | 58.53 | 62.52 | 94.1 |
| chr22 | 66.19 | 47.09 | 98.7 |
| chr2A | 56.56 | 62.86 | 94.6 |
| chr2B | 55.68 | 66.63 | 99.3 |
| chr3 | 54.53 | 65.84 | 98.3 |
| chr4 | 51.96 | 67.09 | 99.9 |
| chr5 | 54.39 | 65.44 | 97.8 |
| chr6 | 55.42 | 66.22 | 98.8 |
| chr7 | 55.12 | 61.26 | 92.6 |
| chr8 | 55.49 | 63.44 | 95.3 |
| chr9 | 57 | 62.46 | 94.1 |
| chrM | 72.64 | 104.49 |  |
| chrX | 52.47 | 61.54 | 92.9 |
| Total | 53.18 | 60.12 |  |
